# Identification of A Disintegrin and Metalloproteinase 9 domain (ADAM9) required in the early stages of encephalomyocarditis virus infection

**DOI:** 10.1101/491068

**Authors:** Lindsey E. Bazzone, Michael King, Christopher R. MacKay, Pyae P. Kyawe, Paul Meraner, Daniel Lindstrom, Joselyn Rojas-Quintero, Caroline A. Owen, Jennifer P. Wang, Abraham L. Brass, Evelyn A. Kurt-Jones, Robert W. Finberg

## Abstract

Encephalomyocarditis virus (EMCV) is a picornavirus that produces lytic infections in murine and human cells. Employing a genome-wide CRISPR-Cas9 knockout screen to find host factors required for EMCV infection, we identified a role for ADAM9 in EMCV infection. CRISPR-mediated deletion of ADAM9 in multiple human cell lines rendered the cells highly resistant to EMCV infection and cell death. Primary fibroblasts from ADAM9 KO mice were also strongly resistant to EMCV infection and cell death. In contrast, ADAM9 KO and WT cells were equally susceptible to infection with other viruses, including the picornavirus Coxsackie virus B. ADAM9 KO cells failed to produce viral progeny when incubated with EMCV. However, bypassing EMCV entry into cells through delivery of viral RNA directly to the cytosol yielded infectious EMCV virions from ADAM9 KO cells, suggesting that ADAM9 is not required for EMCV replication post-entry. These findings establish that ADAM9 is required for the early stage of EMCV infection, likely for virus entry or viral genome delivery to the cytosol.

**Importance:** Viral myocarditis is a leading cause of death in the U.S., contributing to numerous unexplained deaths in people ≤ 35 years old. Enteroviruses contribute to many cases of human myocarditis. Encephalomyocarditis virus (EMCV) infection causes viral myocarditis in rodent models but its receptor requirements have not been fully identified. CRISPR-Cas9 screens can identify host dependency factors essential for EMCV infection and enhance our understanding of key events that follow viral infection, potentially leading to new strategies for preventing viral myocarditis. Using a CRISPR-Cas9 screen, we identified <u>A</u> <u>D</u>isintegrin <u>a</u>nd <u>M</u>etalloproteinase 9 Domain (ADAM9) as a major factor required for the early stages of EMCV infection in both human and murine infection.

## Introduction

Encephalomyocarditis virus (EMCV) is a non-enveloped, single-stranded RNA (ssRNA) virus in the Cardiovirus genus of the Picornavirus family and is known to cause myocarditis, diabetes, and neurologic and reproductive disorders in rodents and non-human primates (1). The virus was first isolated in 1944 from a gibbon that died suddenly from pulmonary edema and myocarditis (2) and later from diseased pigs (3). Since its discovery, EMCV has been isolated globally in an extensive range of animal species (4-7). Rodents, specifically rats, are believed to be the natural reservoir hosts of EMCV, while infection of other animal species may result from occasional cross-species transmission by ingestion of contaminated food, water, or infected carcasses (8-11). EMCV has also emerged as a pathogen capable of causing large zoonotic pandemics and decimating domestic animal populations, making it an important veterinary pathogen. While human infections are rare, EMCV can cause symptomatic disease in humans, manifesting as a mild, non-specific febrile illness (12-15). Infection is more prevalent among humans with occupational exposure to animals, particularly hunters (16-18), suggesting a strong zoonotic potential for EMCV. While serious human EMCV infections are generally rare, EMCV rapidly kills human cells such as HeLa cells as well as primary human cells in culture (19, 20).

EMCV is a well-accepted and widely used model for studying mechanisms of virus-mediated immune suppression, viral myocarditis, and insulin-dependent diabetes (21-25). However, little is known about the receptor requirements of EMCV. The virus receptor on host cells is often a key factor in influencing viral tropism for particular tissues, which subsequently results in various disease manifestations of infection. Thus, understanding viral pathogenesis often hinges on identifying the cellular molecules that the virus binds to facilitate cell entry and subsequent infection. Here, we employed a functional genomics approach to identify genes responsible for EMCV-induced lytic infection in both human and murine cells. Using a genome-wide CRISPR-Cas9 screen, we identified ADAM9 as a major EMCV dependency factor (EDF). ADAMs (<u>A</u> <u>D</u>isintegrin <u>A</u>nd <u>M</u>etalloproteinase domain) are a family of transmembrane metalloproteinases that play important roles in growth factor and cytokine signaling as well as cell-cell signaling, adhesion and extracellular matrix remodeling (26-35). In animals, including humans, ADAM9 is ubiquitously expressed in cells of the developing heart, brain, retina, lung, fibroblasts, neutrophils, and platelets (27, 30, 34-50). Approximately half of the ADAM family members, including ADAM9, have proteolytic capabilities that modulate the activity of cytokines, chemokines, and growth factors, their associated receptors and cell adhesion molecules (27, 35, 37, 45). ADAMs have been implicated in a range of human cancers, inflammatory diseases, wound healing, and microbial infections; however, very little is known about the role of ADAMs in viral infection. This study demonstrates that ADAM9 functions as a major EDF involved in the early infection of both human and murine cells.

## Results

### CRISPR-Cas9 Screening Identifies EMCV Dependency Factors (EDFs)

EMCV infection is rapidly lytic in human and murine cells (51-54). We took advantage of the high lytic potential of EMCV and the power of CRISPR genetic screening (53, 55) to discover virus-host interaction genes that mediated virus infection and, thus, rendered the cells susceptible to EMCV-induced cell death. HeLa cells stably expressing Cas9 were used for screening (53, 55). In initial optimization experiments, we determined that HeLa cells were killed by EMCV within 24 h of infection at a multiplicity of infection (MOI) ≥ 0.1. The rapid lysis of HeLa cells with EMCV infection allowed us to screen for EDFs using pooled single-guide RNAs (sgRNAs) since we could identify such mutant cells by their resistance to EMCV-induced cell death, i.e., these mutants would no longer be susceptible to EMCV infection and would survive EMCV challenge.

We screened for EDFs using CRISPR-Cas9 pooled human gene screen (**Fig. 1**). H1-HeLa cells which stably express a human-codon optimized *S. pyogenes* Cas9 were transduced with GeCKOv2 sgRNA library (53, 54), a complex lentiviral library that targets 19,052 human genes. The GeCKOv2 library contains six unique sgRNAs per gene in two half-libraries (A and B). Libraries A and B each contain three unique sgRNAs per gene and the lentiviral libraries were transduced in parallel into H1-HeLa cells. Stable sgRNA-expressing cells were selected with puromycin for 48 h to allow sgRNA-guided CRISPR-Cas9-mediated gene deletion within each pool. The CRISPR-Cas9-sgRNA transduced H1-HeLa cells were then infected with EMCV ATCC strain VR-129B (MOI = 10). Surviving cells in each pool (<5% of the starting population) were collected 5 d post infection and re-challenged with EMCV and cultured for an additional 7 d (**Fig. 1A**). After two rounds of EMCV selection, genomic DNA was recovered from the surviving cells and sgRNA sequences were amplified from integrated proviruses by PCR and deep sequenced as described (53).

**Figure 1:**
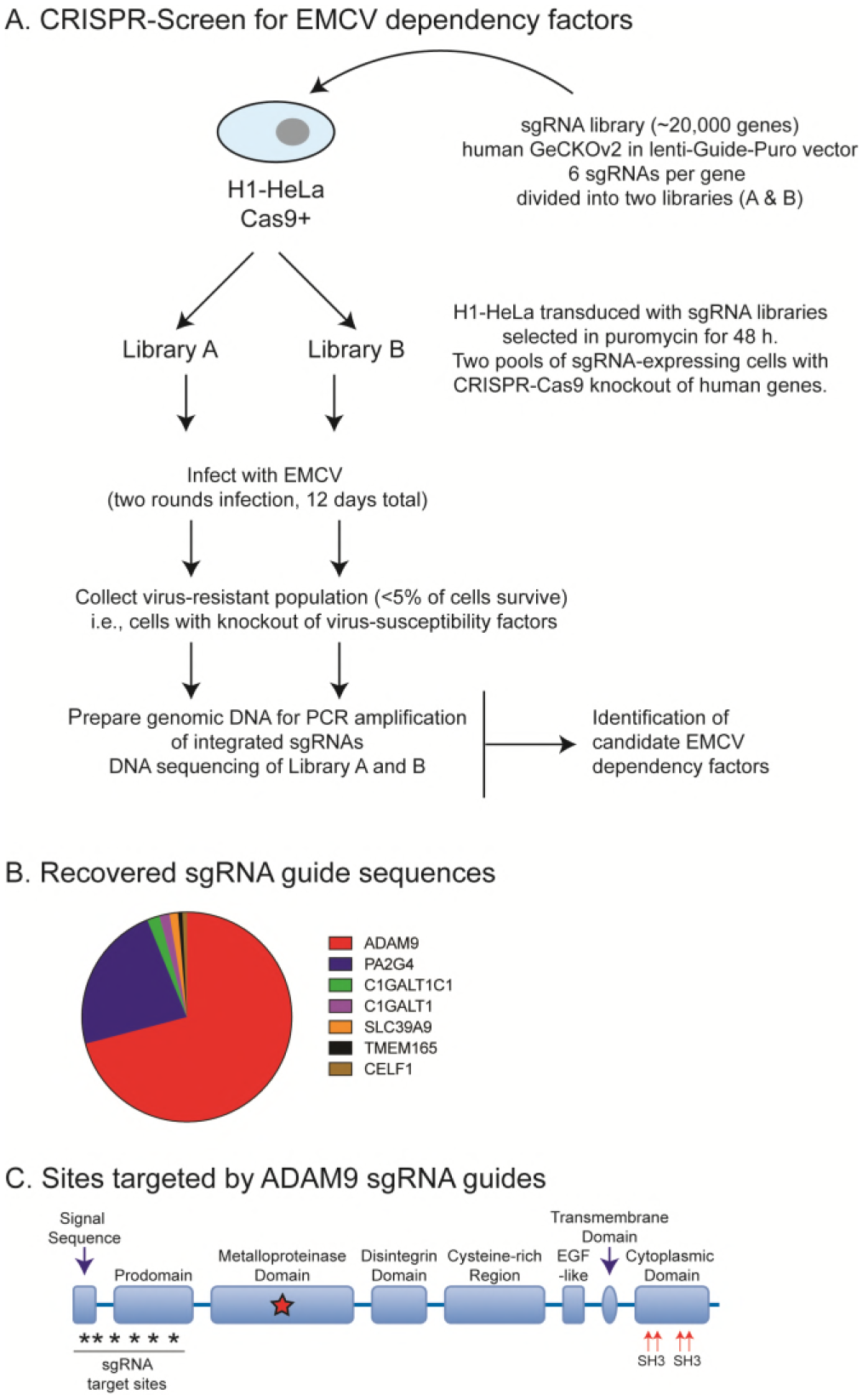
CRISPR-Cas9 knockout screen for EDFs. **(A)** HeLa cells expressing Cas9 were transduced with GeCKOv2 sgRNA library and guide sequences were retrieved from cells that survived multiple rounds of EMCV infection. **(B)** ADAM9 sgRNA sequences were highly represented in recovered clones; 6 of the 6 sgRNAs targeting ADAM9 were present in recovered clones. GeCKOv2 screen results for genes conferring susceptibility to EMCV-induced cell death. Of the pooled CRISPR-targeted genes, ADAM9 and PA2G4 targeted guide sequences accounted for ~70% and ~22%, respectively. **(C)** Domain structure and sites targeted by sgRNAs.

The screen was performed twice and seven candidates (ADAM9, PA2G4, C1GALT1C1, C1GALT1, SLC39A9, TMEM165, and CELF1) were highly enriched in replicate pools (**Fig. 1B**). The top candidate in each pool in replicate experiments was ADAM9, with 6 of 6 sgRNA sequences targeting ADAM9 being significantly enriched (>600 reads) (**Fig. 1B**). ADAM9 sgRNAs were highly enriched in both Library A and Library B pools. Overall, ADAM9 sgRNAs accounted for >70% of the over-represented (>20 reads) sgRNAs recovered from EMCV-resistant cells. The second most abundant candidate was PA2G4, which accounted for >22% of the over-represented sgRNAs. ADAM9 belongs to the ADAM family of type I transmembrane proteins in the zinc-dependent metalloproteinase (MP) superfamily and are comprised of 6 different domains each with distinct functions: 1) a pro-domain that maintains the MP domain in latent form, 2) MP domain, 3) disintegrin domain, which binds integrins to regulate cell-cell or cell-matrix interaction and/or cell migration, 4) a cysteine-rich and epidermal-like growth factor (EGF) domain, thought to promote cell-cell fusion and/or regulate cell adhesion, 5) transmembrane domain, which anchors to the cell surface, and 6) a cytoplasmic tail involved in intracellular signaling (**Fig. 1C**).

### ADAM9 is required for EMCV-induced cell death and virus replication in human and murine cells

To determine the role of ADAM9 in EMCV infection, we used CRISPR-Cas9 and ADAM9-specific sgRNAs to disrupt ADAM9 expression in HeLa and HEK293T human cell lines. WT and ADAM9-deficient cloned cells were tested for their susceptibility to EMCV-induced cell death by assessing cellular ATP levels in a luciferase-based assay as a measure for cell viability. ADAM9 KO clones were resistant to EMCV infection compared to their WT counterparts (**Fig. 2**). WT and ADAM9 KO cells were challenged with EMCV over a range of MOIs and cell viability and EMCV replication were measured after 24 h (**Fig. 2**). WT HeLa cells were sensitive to EMCV-induced cell death at MOI ≥ 0.1 (**Fig. 2A**). In contrast, ADAM9 KO HeLa cells were protected from EMCV-induced cell death even at MOI = 10. At MOI = 1, ADAM9 KO HeLa cells were > 99% viable while < 1% of WT HeLa cells survived EMCV challenge. We also examined EMCV-induced cell death in HEK293T cells (**Fig. 2C**). WT HEK293T cells were very sensitive to EMCV-induced cell death at all MOIs, with ≤ 5% survival of cells at doses as low as MOI = 0.001. In contrast, ADAM9 KO HEK293T cells survived EMCV challenge at doses up to MOI = 10. Thus, loss of ADAM9 expression rendered HEK293T cells >6 logs less sensitive to EMCV-induced cell death compared to WT cells. We confirmed the role of ADAM9 in EMCV infection by measuring virus replication by plaque assays of culture supernatants. Replication of EMCV was markedly reduced in ADAM9 KO compared to WT cells, suggesting that ADAM9 might be required for EMCV binding and/or entry or post-entry transcription of the viral genome (**Fig. 2B,D**).

**Figure 2:**
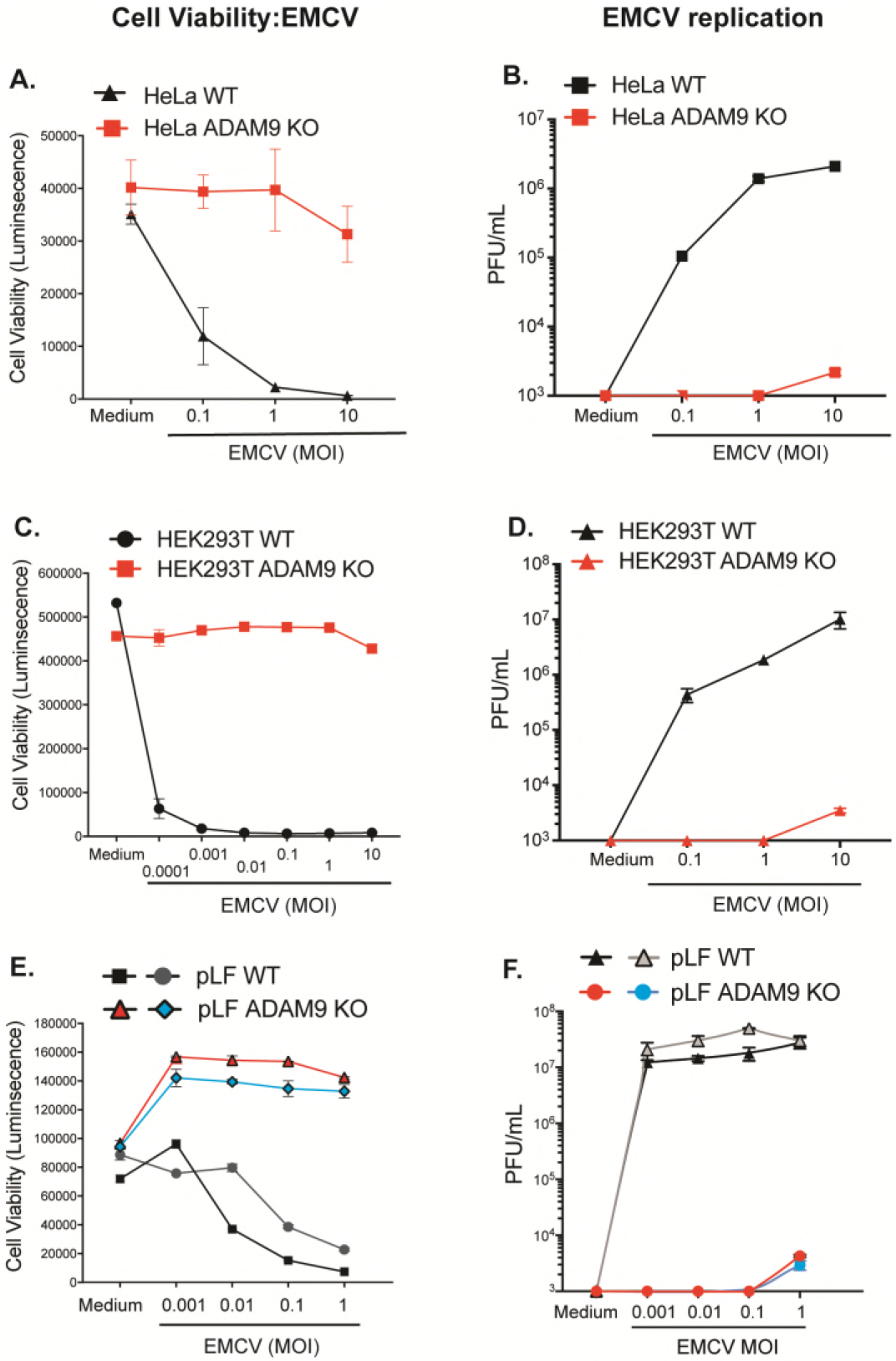
ADAM9 is a major dependency factor for EMCV infection of human and mouse cells. WT and ADAM9 KO HeLa cells **(A,B)** and HEK293T cells **(C,D)**, and two independent sets of primary lung fibroblasts (pLF) isolated from WT and ADAM9 KO mice **(E,F)** were incubated in medium alone or infected with EMCV at varying multiplicities of infection (MOI) for 1 h at 37 °C, washed to remove free virus, and cultured for 24 h at 37 °C. Cell viability: **(A,C,E)** ATP was quantified in the cell monolayers using CellGlo reagent, revealing that ADAM9 KO cells are highly resistant to EMCV-induced cell death. Virus replication: **(B,D,F)** EMCV viral titers, measured on BHK21 monolayers and expressed as plaque forming units (PFU) per mL cell supernatant, showed only low amounts of replicating virus in ADAM9 KO cells. Cell viability data are mean ± SD of 3 replicate wells; plaque assay data are mean ± SD of 2 replicate wells at each dilution. Biological replicate experiments were performed at least twice with similar results.

We next asked if ADAM9 was required for EMCV infection of primary murine cells. Primary lung fibroblasts (pLF) from mice with targeted deletion of ADAM9 (43) and their WT C57BL/6J counterparts were challenged with EMCV. WT pLF were susceptible to EMCV-induced cell death (**Fig. 2E**) and produced high titers of EMCV (**Fig. 2F**). In contrast, ADAM9 KO pLF were highly resistant to EMCV killing and were unable to support EMCV replication. At the highest MOI tested, ADAM9 KO pLF demonstrated some limited EMCV replication, but production of infectious EMCV virions was severely attenuated with an approximate 4-log decrease in infectious virion production (plaque-forming units or PFU/mL) by ADAM9 KO compared to WT cultures. These results suggested that murine ADAM9, like human ADAM9, plays a major role in EMCV-specific lytic infection of cells.

### The requirement for ADAM9 in EMCV-induced cell death is not strain-specific

Previous EMCV studies suggest that different strains of EMCV may bind to different receptors and that some EMCV strains may use multiple pathways (56, 57). In addition, EMCV strains with different tissue tropism have been identified (1). To determine if the requirement for ADAM9 was EMCV strain-specific, we infected WT and ADAM9 KO immortalized mouse lung fibroblasts (iLF) cells with either EMCV (VR129B, ATCC) or with EMCV-M strain (VR-1479, ATCC). ADAM9 KO cells were resistant to both EMCV strains while WT cells were susceptible (**Fig. 3A,B**). We also tested if ADAM9 deficiency had an impact on replication of other RNA and DNA viruses. WT and ADAM9 KO HeLa cells were challenged with Coxsackie virus B3 (CVB3), vesicular stomatitis virus (VSV), influenza A virus (IAV), or herpes simplex virus-1 (HSV-1). WT and ADAM9 knockout cells were equally susceptible to infection with CVB3, VSV, IAV, and HSV-1 (**Fig. 3C,D,E,F**). Cells infected with VSV showed equal yields of infectious VSV from WT and ADAM9 KO pLF (**Fig. 4C**), suggesting that ADAM9 deficiency did not globally impact RNA virus replication.

**Figure 3:**
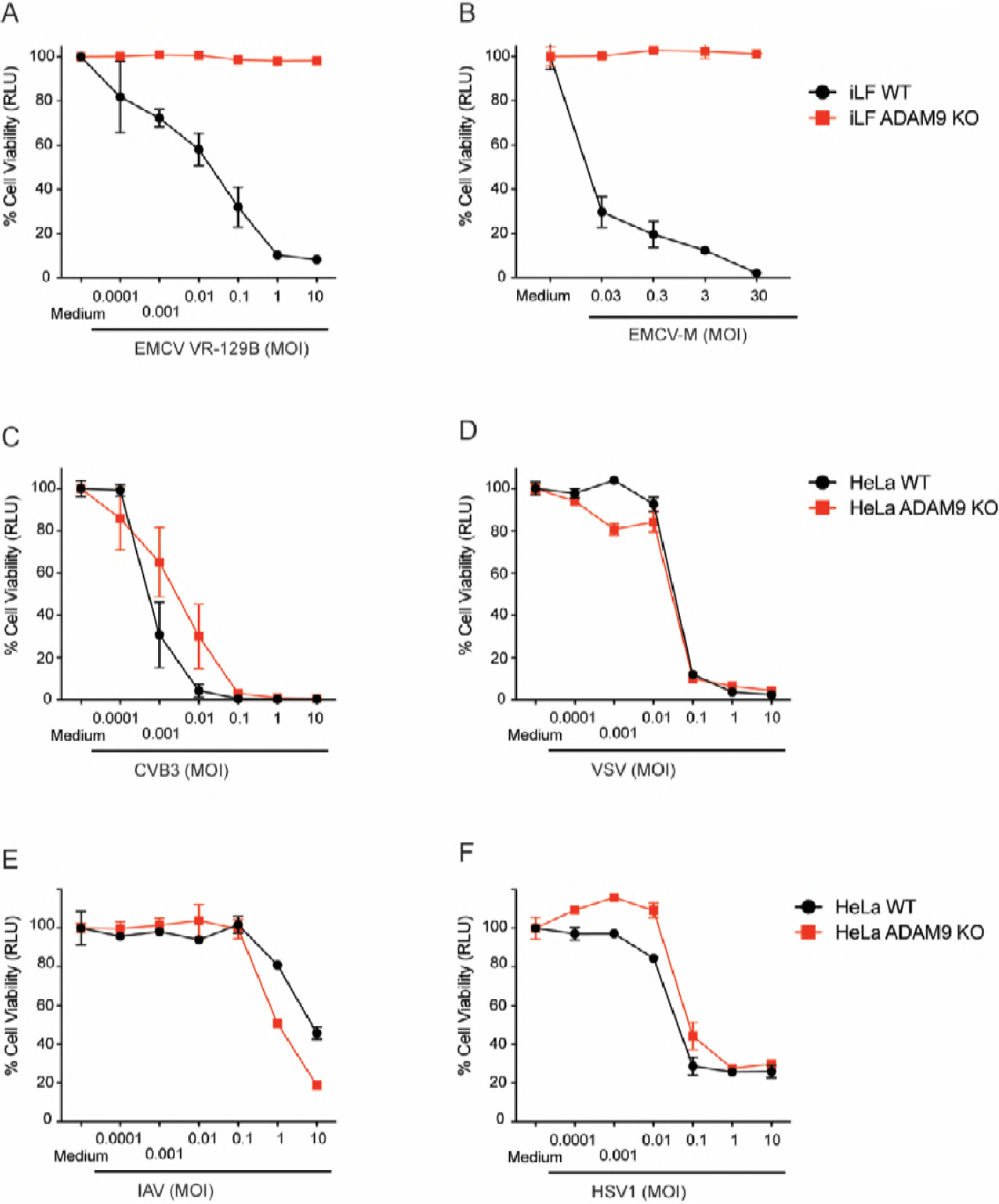
ADAM9 is required for EMCV infection, but does not affect infection by other RNA or DNA viruses. WT and ADAM9 KO cells, including immortalized lung fibroblasts (iLF) and HeLa cells, were challenged with **(A)** EMCV VR-129B, **(B)** M-variant VR-1479, **(C)** Coxsackie virus B3 (CVB3), **(D)** vesicular stomatitis virus (VSV), **(E)** influenza A virus (IAV), or **(F)** herpes simplex virus-1 (HSV-1) at varying MOI or incubated in medium alone for 1 h at 37 °C. Cell monolayers were washed to remove free virus and incubated for 24 h at 37 °C. Viability of infected cells relative to uninfected (medium only) controls was determined with a CellGlo ATP assay and is expressed in relative luciferase units (RLU). Data are mean ± SD of 3 replicate wells. Biological replicate experiments were performed at least twice with similar results.

**Figure 4:**
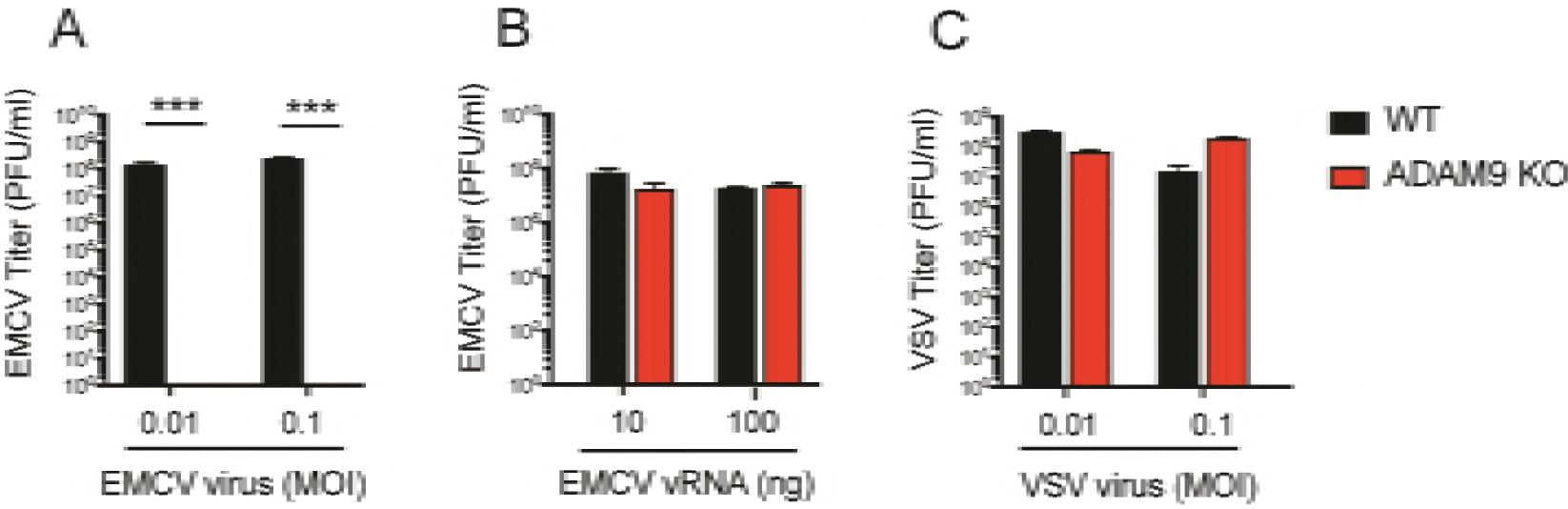
ADAM9 is required for EMCV infection but is not directly required for genome replication in the cytosol. WT and ADAM9 KO immortalized lung fibroblasts (iLF) were **(A)** incubated with EMCV virus for 1 h at 37 °C and washed or **(B)** transfected with EMCV viral RNA (vRNA) in lipofectamine, or **(C)** incubated with VSV. Culture supernatants were harvested after 18 h. Virus replication was measured by quantifying plaque-forming units (PFU) on BHK-21 cells. Data shown are mean ± SD of 3 replicate wells. ***, *P* <0.0001, WT vs ADAM9 KO.

### ADAM9 is essential for EMCV infection, but is not directly involved in post-entry replication of viral genomes in the cytosol

To determine whether ADAM9 is required for virus binding/entry and/or required at a later step in virus replication post-entry, we transfected infectious EMCV viral RNA (vRNA) into iLF using lipofectamine, thus by-passing the requirement for receptor-dependent entry events. iLFs were either infected with intact EMCV virions for 1 h at 37°C, washed to remove residual free virus, and cultured for 18 h at 37 °C (**Fig. 4A**) or transfected with EMCV vRNA in lipofectamine and incubated at 37 °C (**Fig. 4B**). iLFs were also infected with VSV, as a positive control, for 1 h at 37 °C, washed, and cultured (**Fig. 4C**). After 18 h, culture supernatants were harvested and EMCV, or VSV, replication was determined by quantifying infectious virions in culture supernatants by plaque assay. EMCV infection of WT cells resulted in productive infection of cells (at least 8 logs within 18 h), while infection of ADAM KO cells, resulted in no virus production (**Fig 4A**). When receptor-mediated cell entry is bypassed by transfection of infectious vRNA directly into the cell, EMCV is capable of replicating in both WT and ADAM9 KO cells (**Fig. 4B**). Together, these results demonstrate that ADAM9 is not directly involved in EMCV replication steps post-entry, but rather facilitates EMCV entry and/or genome delivery into the cell where EMCV vRNA genomes are replicated in an ADAM9-independent manner.

### Rescue of ADAM9 in KO cells restores susceptibility to EMCV

We next examined if rescue of ADAM9 expression to ADAM9 KO cells restores EMCV susceptibility. ADAM9 KO and WT HeLa and mouse iLF cells, were transduced with retroviral vectors to express murine ADAM9 (mADAM9) or GFP as a control. Transduced cells were selected and cloned by limiting dilution. Expression of ADAM9 was confirmed by western blot analysis (**Fig. 5A**). ADAM9 KO cells rescued with intact wild-type (WT) mADAM9 became susceptible to EMCV-induced cell death (**Fig. 5B**), demonstrating that ADAM9 is both necessary and sufficient to confer EMCV susceptibility.

**Figure 5:**
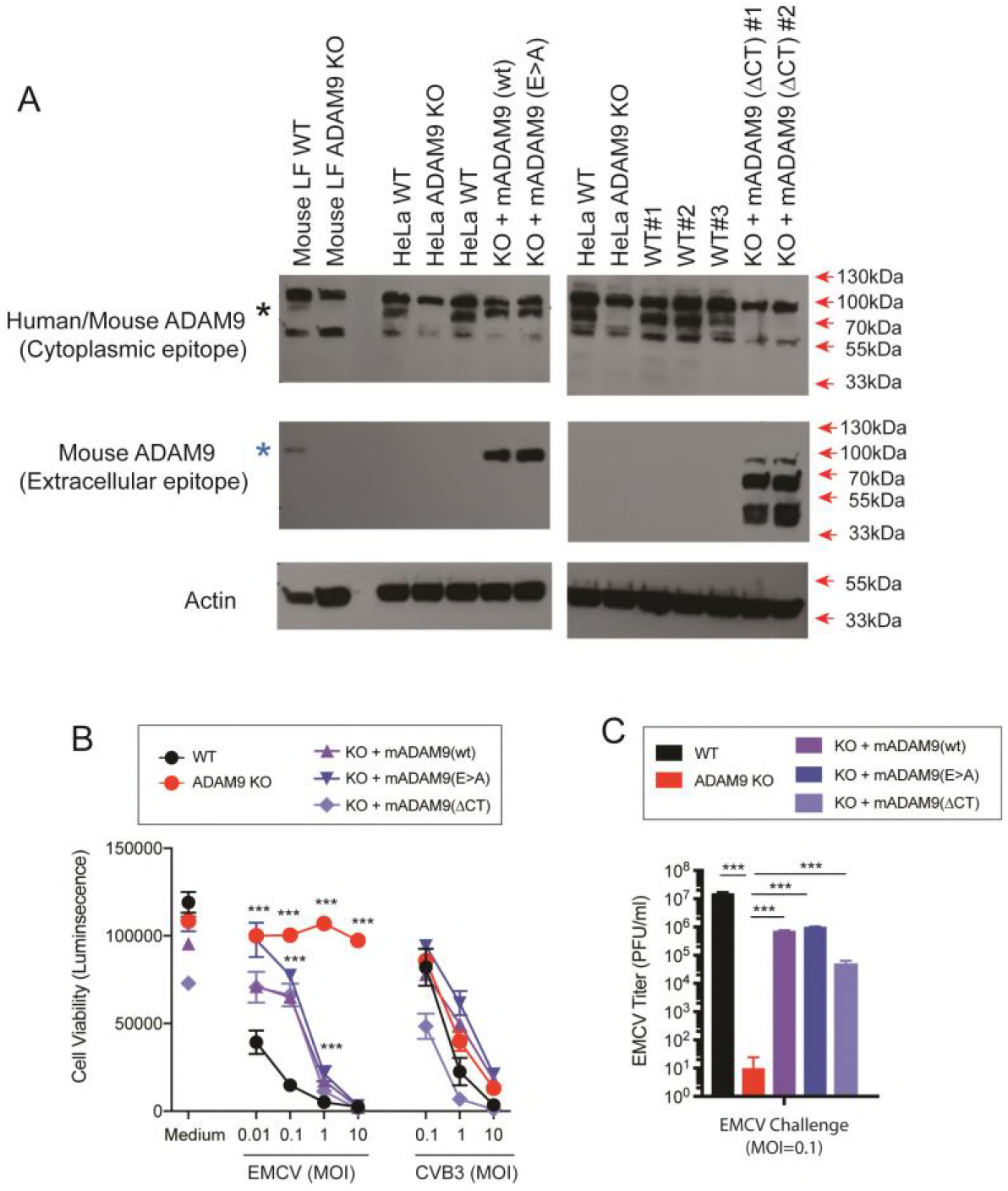
Restoration of ADAM9 expression restores EMCV replication in ADAM9 KO cells. WT and ADAM9 KO HeLa cells were transduced with retroviral vectors with wild-type (WT) murine ADAM9 (mADAM9), catalytically inactive mutant ADAM9 (E>A), cytoplasmic tail deleted (ΔCT) ADAM9 constructs, or GFP control vectors. Transduced cells were selected and cloned by limiting dilution, and expression of mADAM9 was confirmed by western blot analysis. **(A)** WT, KO and rescue cell lysates were prepared in RIPA buffer and run on 12% SDS-PAGE gels and transferred to PVDF membranes. Membranes were blocked with 3% BSA and incubated with ADAM9 antibodies. Upper panel, rabbit anti-human ADAM9 (Cell signaling #2099) that detects an epitope in the intracellular domain of human ADAM9 and cross-reacts with mouse ADAM9. Middle panel, goat anti-mouse ADAM9 (R&D systems AF949) that detects the extracellular domain of murine ADAM9, but not human ADAM9. Lower panel, anti-Actin (Santa Cruz sc-1616) which detects both human and murine ϟ-Actin. Bands were visualized using HRP and ECL reagent. Upper panel: WT but not KO cells expressed human ADAM9. Middle panel: Rescue but not KO cells expressed murine ADAM9. Lower panel: Actin loading control. **(B)** WT clones, ADAM9 KO clones and rescued ADAM9-expressing clones were infected with EMCV or CVB3 at varying MOI and incubated at 37 °C for 24 h. Viability of EMCV-infected and CVB3-infected clones was measured by CellGlo ATP luminescence. **(C)** EMCV replication was quantified in infected culture supernatants by plaque assay using BHK-21 cells. Neither the functional sequence of the ADAM9 metalloproteinase domain nor the cytoplasmic tail are required for EMCV infection. ***, *P* <0.0001, KO vs. WT; KO vs. rescue.

ADAM9 is a multi-functional protein and has several domains that participate in a variety of cell processes. ADAM9 controls receptor-mediated signaling via the proteolytic function of its MP domain and signals via its SH3 domains in the cytoplasmic tail. To determine if mutating the MP domain of ADAM9 impacts susceptibility to EMCV, we rescued ADAM9 KO cells with an enzyme-dead mutant ADAM9 containing a Glu to Ala (E>A) substitution in the MP active site (58). The MP mutant ADAM9 restored EMCV susceptibility to KO cells (**Fig. 5B,C**). Thus, EMCV susceptibility in KO cells rescue with (E>A) mutant was indistinguishable from rescue with WT ADAM9 indicating that the active MP domain is not required for EMCV entry into cells. We next asked if the intracellular signaling domain of ADAM9 was necessary to render cells susceptible to EMCV. We replaced the cytosolic domain of ADAM9 with a V5-epitope tag and expressed the cytoplasmic tail deleted (ΔCT) mutant in KO cells (**Fig. 5B,C**). EMCV susceptibility was restored with ADAM9-ΔCT rescue suggesting that the cytoplasmic tail of ADAM9 does not play a role in EMCV susceptibility.

We also challenged WT, ADAM9 KO and rescued KO cells with varying doses of CVB3. As shown in **Fig. 5B**, we did not observe any differences in susceptibility to CVB3 infection in WT, KO and rescued cells, confirming that ADAM9 is specifically required for EMCV infection.

## Discussion

Forward genetic screens allow an unbiased and comprehensive approach to discovering host factors that promote or restrict virus replication. Recent advances in CRISPR-Cas9 have allowed highly efficient generation of knockout cells with marked phenotypes for the identification of virus dependency factors with very few false positive hits. CRISPR-Cas9 screens using multiplexed pools of sgRNAs that cover the entire human genome have proved to be a powerful technologic advance and have been successfully used to identify receptors and other cellular dependency factors for several viruses, including influenza A virus, Zika virus, West Nile virus, dengue virus and hepatitis C virus (53, 54, 59-61), reviewed in (55).

In the present study, we employed a genome-wide CRISPR-Cas9 screen to identify EDFs. The screen recovered multiple host factors, including those involved in the post-translational modification of cell surface proteins (C1GALT1C1, C1GALT1, TMEM165), cellular proliferation (PA2G4), the regulation of mRNA processing (CELF1), and the metalloproteinase ADAM9. We focused on ADAM9 because it was the most strongly enriched candidate in the screen and scored with multiple orthologous sgRNAs. Using a diverse panel of CRISPR-engineered ADAM9 KO human cell lines, as well as primary and immortalized murine cell lines from ADAM9 KO mice, we confirmed ADAM9’s involvement in EMCV infection and demonstrated that loss of ADAM9 results in a profound block to the early stages of EMCV infection. To our knowledge, this study is the first to define a direct role for ADAM proteins in EMCV infection.

Previous studies have suggested different proteins that may be involved in susceptibility to EMCV, but definitive proof (including reconstitution of knockout cells) has not been forthcoming. For example, vascular cell adhesion molecule 1 (VCAM-1), a member of the immunoglobulin superfamily, found on primary vascular endothelial cells has previously been reported as a receptor for EMC-D strain (62). However, cells lacking VCAM-1 have also been shown to be susceptible to EMCV infection via a yet uncharacterized 70 kDa glycoprotein hypothesized to be responsible for mediating virus binding and attachment in these cells (56). Infection of human cardiomyocytes with two different strains of EMCV demonstrated a requirement for sialic acid and heparan sulfate in one EMCV strain but not the other, raising the possibility that different EMCV strains may bind to different polysaccharides in order to initiate infection (57). While different molecules have been implicated as receptors or attachment factors for specific strains of EMCV, the provenance of these strains has been difficult to authenticate, and we have not been able to obtain many of the strains previously reported to have specific receptor or attachment requirements. However, in this study, we were able to confirm that the two ATCC strains of EMCV both use ADAM9 as their major dependency factor. ADAM9 is likely the 70 kDa putative EMCV receptor identified by Jin et al. (56). Furthermore, we showed that the dependency on ADAM9 for infection is unique to EMCV as infection with other RNA and DNA viruses was not affected by ADAM9 deficiency. Thus, although previous studies have identified certain cell surface molecules as potential EMCV receptors or EDFs, this is the first report of a definitive, widely-expressed cell surface protein utilized by multiple EMCV strains that is critical for EMCV infection. Future experiments will be necessary to define the role of ADAM9 and other cell surface proteins as binding or entry receptors or receptor co-factors for EMCV variant strains, as the use of alternative receptors or attachment proteins may impact tissue tropism and ultimately EMCV pathogenesis.

ADAMs have been shown to regulate receptor signaling through a variety of mechanisms many of which involve the catalytically active MP domain (27, 28, 30, 31, 33, 35, 42, 63-65). This raises the possibility that ADAM9 may function as a cofactor for EMCV infection. Given the multifunctional domains of ADAMs, it is also possible that ADAM9 plays a role in the immune response to EMCV infection following virus entry, for example by delivering the viral genome to intracellular innate immune sensors triggering an appropriate antiviral response. Alternatively, it is also possible that ADAM9 may be involved in innate immune evasion by EMCV, such as allowing EMCV to enter the cell undetected or by skewing the innate antiviral signaling pathways. Of ADAM9’s multiple domains, we initially hypothesized that the enzymatically active MP domain and the intracellular cytoplasmic domain containing SH3 binding domains might be involved in EMCV infection. However, our rescue experiments showed that neither the metalloproteinase activity nor the cytoplasmic tail of ADAM9 is necessary to support lytic EMCV infection. Furthermore, the finding that EMCV infection does not require a catalytically active proteinase domain suggests that ADAM9 may not act as a proteolytic cofactor and further supports a direct virus-receptor interaction between ADAM9 and EMCV or a non-enzymatic protein-protein interaction of ADAM9 with the EMCV receptor that is required to facilitate virus entry.

Our data support that ADAM9 is a crucial protein for EMCV infection; specifically, ADAM9 knockout cells from two mammalian species are not infected by two different EMCV strains, reconstitution of ADAM9 permits EMCV infection, and virus replication is unaffected by ADAM9 deficiency when viral RNA is transfected directly into the cytosol. EMCV may directly bind to ADAM9, but we and others (57) have observed polysaccharide-related binding that may facilitate productive infection. Also, as a disintegrin, ADAM9 is known to affect expression of other cell surface proteins (27, 38), thus expression of ADAM9 could lead to uncovering of another protein that is the actual binding “receptor” protein. Since various domains and functions of ADAM9 have been identified, future experiments will yield a fuller definition of how ADAM9 facilitates entry of EMCV into the cell.

In addition to ADAM9, our CRISPR-Cas9 screen in HeLa cells also identified another gene, *PA2G4*, which is likely also involved in EMCV susceptibility (**Fig. 1**). The *PA2G4* gene encodes proliferation-associated protein 2G4, also known as ErbB3-binding protein 1 (EBP1), which plays a role in cell signal transduction, transcription and translation, and has been shown to suppress growth in certain cancers, including breast and prostate cancers (66-72). How EBP-1 affects EMCV susceptibility and its relationship to ADAM9 is currently under investigation.

The discovery of ADAM9 as a surface protein required for EMCV infection has significant implications on our understanding of virus-host interactions and downstream effects of EMCV infection. We are defining ADAM9’s precise role in downstream aspects of EMCV infection. Detailed insight into the mechanism of the EMCV-ADAM9 interaction is crucial to understanding viral pathogenesis. Functional studies examining mechanisms of EMCV binding and entry and potential for ADAM9-mediated cell signaling and cytokine production in response to EMCV infection are currently being explored.

## Materials and Methods

### GeCKOv2 library screen

H1-HeLa cells expressing the GeCKOv2 CRISPR library (Agilent) were plated for full coverage of the library (both pools A and B). These cells were challenged with EMCV at MOI = 10 until < 5% of living cells remained. The cells were washed and re-plated for another challenge of EMCV for confirmation of resistance. gDNA was isolated by expanding the remaining cells and using Qiagen DNeasy Blood and Tissue kit on 2 × 10^6^ cells. PCR of was performed with lentiGP-1_F and lentiGP-3_R (sequences below) with the product spanning the sgRNA guide sequences. This PCR product was processed and sequenced using the Illumina NextSeq 500. The library screen was performed twice for each pool of the GeCKOv2 library.

### Cell viability assay

Cells were plated at 5,000 cells per well in duplicate on a 96-well flat-bottomed plate and incubated overnight at 37 °C. The cells were challenged with virus at the respective MOI for 1 h, washed with PBS; and 150 μL of fresh complete media was added to each well. The plates were incubated for 24 h post infection at 37 °C, 10% CO_2_. The media supernatants were collected and 50 μL of fresh media and an equal volume of room temperature CellTiter Glo reagent (Promega G7571) were added to each well and mixed for 2 min at 300 rpm. The reaction was allowed to equilibrate for 10 min, then 80 μL of the reaction transferred to an opaque 96-well plates and the signal was detected by luminometer.

### CRISPR KO

ADAM9 KO lines were created using CRISPR technology. HeLa cells (H1 cell line) and 293T cells with stable expression of Cas9 (53) were transiently transfected with ADAM9
CRISPR Guide RNA #1 (CACTGACCATCCCAATATAC) and #3 (CTTATGAAATTATAACTCCT) from Genscript sgRNA guide sequences were based on the GeCKO Library designed by Feng Zhang’s laboratory at the Broad Institute, see http://genome-engineering.org/gecko/). Cells were incubated for 3 d at 37 °C to allow Cas9-mediated gene deletion. Cells were screened by limiting dilution cloning. Knockout clones were validated by western blotting.

### ADAM9 constructs

All mouse ADAM9 constructs were made using a Gibson assembly strategy. The pQCXIP vector (53) was cut with Not I restriction enzyme. Two-step PCR of pcDNA-mouse ADAM9 WT and E>A for full length ADAM9 (Blobel lab) (mAdam9FL-F; mAdam9FL-R primer sequences below) was done using Phusion High Fidelity Master Mix (New England Biolabs M0531) as recommended by the manufacturer. The template for the PCR for the ΔCT mutant was the pcDNA mouse ADAM9 WT plasmid (mAdam9FL-F; mAdam9dCT-R primer sequences below) where the reverse primer added a V5 tag as above, and a V5-pQCXIP Gibson adaptor (V5-pQCXIP sense primer sequence below) added the stop codon to the end of the transmembrane domain, replacing the intracellular domain. The PCR products were purified (Qiagen). Gene assembly was done with Gibson Assembly master mix (New England Biolabs E2611) per instructions of the manufacturer into the pQCXIP retroviral backbone. Stbl3™ One Shot™ competent cells (ThermoFisher) were transformed with the Gibson assembly product. Minipreps were made from single colonies of transformed bacteria and sequenced. Maxipreps of the sequenced plasmids were made and purified using the Qiagen HiSpeed Maxi Kit.

### Rescue cell lines

ADAM9 rescue of the knockout cell lines was done by transduction. Retrovirus was made using 293FT cells transfected with retroviral transfer constructs with a pQCXIP backbone (GFP ‒ Addgene #73014; mouse Adam9 WT; mouse Adam9 E>A; mouse Adam9 ΔCytoplasmic tail), pMD 2.G, and VSV-G packaging plasmid using TransIT-293 transfection reagent (Mirus 2704) per protocol (53). After 24 h the media was replaced with complete DMEM. At 24 h and 48 h supernatant was collected (and replaced with complete DMEM as needed) and filtered through 0.45 μm filter. Virus supernatants were used immediately or frozen at −80 °C in single use aliquots.

Cell lines to be rescued were plated on 6-well plates and incubated overnight prior to retroviral transduction. Culture medium was replaced with fresh complete DMEM and retroviral supernatant was added dropwise to wells. Cells were incubated at 37 °C for 2-3 d. Transduced cells were cloned by limiting dilution into 96-well plates and grown at 37 °C. Cloned mouse ADAM9 rescue cell lines were confirmed by western blot and genomic DNA sequencing.

### Isolation of mouse pLFs and backcrossing of KO mice

All procedures conducted on mice were approved by the Brigham and Women’s Hospital Institutional Animal Care and Use Committee. Lung fibroblasts were isolated from C57BL/6 WT and ADAM9 KO mice that were in a pure C57BL/6 background. Primary lung fibroblasts were generated from 6-10 week old mice according to a protocol adapted from Yamamoto et al. (73). Euthanized mice were washed with 90% ethanol to reduce contamination. Whole lung lobes were dissected out and place into sterile PBS. Lungs were then submerged in 70% ethanol for 20 seconds, and then placed into digestion media (DMEM with 0.25% trypsin) and cut into small pieces using a sterile scalpel. The lung pieces were then transferred into 10 mL of digestion media and incubated with shaking at 37 °C. After 30 min, 5 mL of cold complete DMEM (DMEM, 10% heat-inactivated FBS, 2 mM L-glutamine, 100 U/mL penicillin, 100 μg/mL streptomycin, 250 ng/mL amphotericin B) was added, and the cells were centrifuged for 5 min at 350 × g at 4 °C. The cell pellet was removed and resuspended in fresh complete DMEM. For each mouse, the lung pieces and cells were resuspended in 50 mL of complete DMEM and then plated in a 225 cm^2^ tissue culture flask. The cells were incubated for 10 to 15 days with minimal disruption, with fresh media added after 7 d. For the first passage, the cells and lung pieces were detached from the flasks by scraping, and separated into cells and lung pieces with 70 μm cell strainer. The single-cell suspension was either passaged one or two times when 90% confluent in complete DMEM, or used immediately for stimulation. Fibroblasts were immortalized with SV40.

### Viruses

EMCV VR129B, EMCV-M VR-1479, and VSV strain Indiana were obtained from the American Type Culture Collection (ATCC). IAV WSN/33 was obtained from Dr. Brass (74). HSV-1 KOS was a gift from David Knipe (Harvard Medical School). CVB3 Nancy stocks were from Robert Finberg.

### Preparation of EMCV vRNA

EMCV vRNA was isolated from ultra-pure EMCV stocks using QIAamp Viral RNA Mini Kit (Qiagen) according to the manufacturer’s protocol. Viral RNA concentration was determined by NanoDrop and vRNA stocks were stored at −20 °C until use. Cells were transfected with viral RNA at 100 ng/well with 0.5 μl of Lipofectamine 2000 (ThermoFisher scientific #11668027). After 1 h of transfection, the cells were washed, trypsinized and diluted into BHK-21 cells as described above.

### Plaque Assay

BHK-21 cells were plated on 12-well plates at 7 × 10^5^ cells/well and incubated overnight at 37 °C to allow formation of intact monolayers. Cell growth media was removed and replaced with 450 μL per well of fresh complete DMEM with 10% FBS. Viral supernatant samples were thawed on ice and 10-fold serial dilutions of virus were performed in serum-free DMEM. Diluted supernatants were added to intact BHK-21 monolayers in duplicate at 50 μL per well and mixed by gentle plate shaking (final dilutions between 10^−2^ to 10^−7^). Cells were incubated at 37 °C for 1 h, with gentle shaking every 20 min. After 1 h, virus dilutions were removed by aspiration and cell monolayers were washed with 1X PBS to remove unbound virus. An agarose overlay was added to each well and allowed to solidify at room temperature. Plates were then incubated at 37 °C for 24 h. Cells were then fixed/stained by adding Crystal Violet dissolved in 4% paraformaldehyde (0.5% Crystal Violet (Sigma), 4% paraformaldehyde (Sigma) in PBS (Corning) at room temperature for at least 2 h. Overlays were removed and plaques counted. PFU/mL was calculated by averaging the number of plaques in the duplicate wells, dividing by the dilution factor, and volume applied to the assay wells.

### Primers

lentiGP-1_F 5’AATGGACTATCATATGCTTACCGTAACTTGAAAGTATTTCG 3’

lentiGP-3_R 5’ATGAATACTGCCATTTGTCTCAAGATCTAGTTACGC 3’

mAdam9FL-F 5’CGCTGCAGGAATTGATCCGCGGATCCCGGGCTGCAGGAATTC 3’

mAdam9FL-R 5’CGAGGCCTACCGGTGCGGCCCGAGCGGCCGCCAGTGTGATGGAT 3’

mAdam9ΔCT-R 5’TAATCGTCCCCCTTGTTGCGGCTGCCATTTTCCTCTTTATCAAG**GGTAAGCCTATCCCTAACCCTCTCCTCGGTCTCGATTCTACG** 3’(V5 tag in bold)

V5-pQCXIP (sense) 5’CCCTCTCCTCGGTCTCGATTCTACGTGATAGTAGGCACCGGTAGGCCTCG TACGC 3’

### Western Blot

Following trypsinization, cells were lysed with RIPA buffer with protease inhibitor cocktail (Sigma P8340) incubated at room temperature for 10 min. The lysate was cleared by centrifugation at 15,000 rpm for 10 min to remove cell debris. The concentration of the lysates was determined by BCA Assay (ThermoFisher PI23227). 12% precast gels (Biorad 4561044) were loaded equally with lysate and run at 80 V. The proteins were transferred to a PVDF membrane (Biorad 1620219) with a semi-dry transfer system (Biorad 1703940) for 60 min at 25 V (300 mA max.) The membrane was blocked with 3% BSA or 5% non-fat dry milk in Tris-buffered saline with 0.1% Tween 20 (TBS-T) for at least 1 h. Incubation of the membrane with primary antibodies was done overnight at 4 °C with rocking/shaking. Membranes were washed three times with TBS-T. Secondary HRP-conjugated antibodies were diluted in TBS-T, added to membrane, and incubated for 1-2 h at room temperature with shaking. Membranes were washed five times with TBS-T and visualized with ECL (Thermo Scientific 34080) and exposed on film as needed. To determine loading, the membrane was stripped with Restore Plus Western stripping buffer (Thermo Scientific 46430) for 10 min and washed twice with TBS-T. The membrane was blocked again in 3% BSA in TBS-T for at least 1 h. Anti-actin HRP conjugate was added in TBS-T and incubated for ~1 h at room temperature with shaking. The membrane was then washed five times with TBS-T and visualized as above. Antibodies used were goat anti-mouse ADAM9 (R&D systems AF949 1:1000 dilution, 3% BSA blocking buffer), rabbit anti-human ADAM9 (Cell Signaling #2099 1:1000 dilution, 5% milk blocking buffer), anti-actin HRP conjugate (Santa Cruz sc-1616 1:2000 dilution. 3% BSA blocking buffer), anti-goat IgG HRP (1:2000 dilution, R&D Systems HAF109), and anti-rabbit IgG HRP (1:5000 dilution Vector Labs PI1000).

### Statistical Analysis

Differences in the means of data were compared using unpaired, two-tailed Student’s t-test. ANOVA was used when comparing multiple values. Significant differences were assumed for *P* values < 0.05. All statistical analyses were performed using GraphPad Prism software (version 7.0d; GraphPad, San Diego, CA).

## Funding sources

American Heart Association-Myocarditis Foundation (18POST34030152) to L.E.B.; NIH NIAID (T32 AI095213) to C.R.M.; NIH NIAID (R01 AI116920) to J.P.W.; Burroughs Wellcome Foundation, Bill and Melinda Gates Foundation, Gilead Sciences Inc., NIAID (R01 AI091786) to A.L.B.; NIH NIAID (R01 AI111475), Flight Attendants Medical Research Institute grant (CIA123046), The Brigham and Women’s Hospital–Lovelace Respiratory Research Institute Consortium, and Department of Defense (Congressionally Directed Medical Research Programs) grant (PR152060) to C.A.O.

## Acknowledgments

We thank Melanie Trombly for assistance with manuscript preparation. We also thank Dr. Carl P. Blobel and Dr. Gisela Weskamp for generously providing the plasmids for the wild-type and E>A mutation and the ADAM9 KO mice. Author contributions are as follows: J.P.W., A.L.B., E.A.K.-J., and R.W.F. designed the research; L.E.B, M.K., C.R.M, P.P.K, P.M., and D.L. performed the research; G.W, C.P.B, J.R.-Q., and C.A.O. contributed new reagents/analytic tools and contributed to data interpretation; L.E.B., J.P.W., E.A.K.-J., A.L.B. and R.W.F. contributed to the analysis and wrote the paper. We have no conflicts of interest to report.

